# Oligodendrocyte Precursor Cells Are Co-Opted by the Immune System to Cross-Present Antigen and Mediate Cytotoxicity

**DOI:** 10.1101/461434

**Authors:** Leslie Kirby, Jing Jin, Jaime Gonzalez Cardona, Matthew D. Smith, Kyle A. Martin, Jingya Wang, Hayley Strasburger, Leyla Herbst, Maya Alexis, Jodi Karnell, Todd Davidson, Ranjan Dutta, Joan Goverman, Dwight Bergles, Peter A. Calabresi

## Abstract

Oligodendrocyte precursor cells (OPCs) are abundant in the adult CNS and can be recruited to form new oligodendrocytes and myelin in response to injury or disease. However, in multiple sclerosis (MS), oligodendrocyte regeneration and remyelination are often incomplete, suggesting that recruitment and maturation of OPCs is impaired. MS and the rodent model experimental autoimmune encephalomyelitis (EAE) are characterized by infiltration of activated T-cells into the CNS. To investigate the mechanisms by which this neuroinflammatory process influences OPC mobilization, we performed in vivo fate tracing in an inflammatory demyelinating animal model. Results of our studies showed that the OPC differentiation and myelin production are inhibited by either adoptive transfer of CNS infiltrating cytokine producing effector T-cells or CNS production of interferon gamma (IFNγ), using an astrocyte specific IFNγ transgene model. In both systems, IFNγ changes the profile of OPCs by inducing functional expression of the immunoproteasome and upregulation of MHC class I. OPCs exposed to IFNγ are shown to cross present exogenous antigen to cytotoxic CD8 T-cells, which then produce proteases and FasL that results in subsequent caspase 3/7 activation and OPC death, both in vitro and in vivo. Cross presentation by OPCs is dependent on the cytosolic processing pathway and can be inhibited by small molecules targeting MHC class I antigen processing and the immunoproteasome subunits. Finally, the immunoproteasome subunit, PSMB8, is shown to be markedly increased on Sox10^+^ oligodendrocyte lineage cells only in the demyelinated white matter lesions from patients with MS. These findings support the notion that OPCs have multiple functions beyond differentiation into myelinating cells and adapt to their microenvironment by responding to local cues. In MS, OPCs may be co-opted by the immune system to perpetuate the autoimmune response. Strategies aimed at inhibiting the aberrant immune activation pathways in OPCs may allow more efficient remyelination in MS.

## Introduction

Oligodendrocyte precursor cells (OPCs) that express the proteoglycan neuron-glial antigen 2 (NG2) are a highly dynamic, proliferative group of progenitors that remain abundant in the adult CNS. Differentiation of OPCs into oligodendrocytes allows oligodendrocyte generation to continue into adulthood for adaptive myelination and the ability to regenerate myelin following injury or disease. However, the abundance and tiling-like coverage of NG2^+^ OPCs in both gray and white matter of the CNS, suggests that they may have other functions; indeed, OPCs survey their microenvironment through constant filopodia extension^1–4^, migrate to sites of injury and respond to inflammatory cues^5–8^, behaviors remarkably similar to microglial cells. The significance of these non-progenitor behaviors in both physiological and pathological conditions are not well understood.

In the setting of chronic inflammatory demyelinating diseases such as multiple sclerosis (MS) the process of endogenous remyelination is inefficient, rendering axons susceptible to degeneration through loss of trophic support and direct toxic effects of immune cells. Inflammatory cells and factors participating in MS disease pathology have to be taken into consideration in order to understand the mechanisms underlying remyelination failure. Recently, subsets of immune cells have been shown to have differing roles in response to tissue injury. T regulatory cells (Treg)^7^ and alternatively activated monocytes (M2)^9^ can facilitate endogenous remyelination through direct actions on OPCs, whereas CD4^+^ T-cells of the Th1 and Th17 lineage have been shown to hinder these processes^10–13^. In autoimmune diseases, effector T-lymphocytes have long been understood to be key in the establishment and perpetuation of MS. Both CD4^+^ and CD8^+^ T-cells have been identified in acute and chronic lesions. While MHC class II alleles have been strongly associated with MS risk, CD8^+^ cells still far outnumber CD4^+^ cells in MS lesions^14–16^. In addition, axonal damage closely correlates with CD8^+^ cell number prevalence rather than CD4 prevalence^17^. While CD8^+^ cells exhibit the effector function of contact-mediated cytotoxicity, both CD4^+^ and CD8^+^ cell mediate pathology through cytokine production. The additional function of cytotoxicity provides CD8^+^ cells with a tissue destructive function that has been proposed to be important for MS pathogenesis^18–23^. Importantly, recruitment of OPCs to the demyelinated lesion temporally and spatially overlaps with the persistence of CD4^+^ and CD8^+^ T-cells^24^.

Detection of tissue injury is an important function for the OPC pool, aiding their ability to mobilize an appropriate repair response. Several lines of evidence suggest that OPCs express cytokine receptors, and their differentiation is inhibited by interferon gamma (IFNγ)^25–30^, interleukin 17 (IL-17)^5,6^, and high doses of tumor necrosis factor (TNF)^31,32^, but the mechanistic pathways mediating these effects remain incompletely understood. We and others have previously used *in vivo* genetic fate tracing to track the differentiation of OPCs into mature myelin-producing oligodendrocytes during remyelination in the cuprizone model^33,34^. We also demonstrated that adoptive transfer of myelin-reactive T effector cells inhibits endogenous remyelination in the corpus callosum without causing irreparable damage to the axons, as is common in the spinal cord of animals with experimental autoimmune encephalomyelitis (EAE)^35^.

Based on these prior studies and because the effects in our model system were seen early before secondary inhibitory mechanisms could occur, we hypothesized that inflammatory cytokines might be directly signaling to OPCs and inhibiting their differentiation. Furthermore, the OPC behavior of microenvironment surveillance in both gray and white matter regions begs the question as to other important functions that the OPCs may possess, especially in the context of inflammation. Herein, we show a detailed mechanistic signaling pathway in OPCs that not only diverts them from differentiating into mature oligodendrocytes but also induces expression of antigen-presenting capacity through induction of the immunoproteasome. OPCs exposed to inflammatory cytokine, IFNγ, highly express MHC class I molecules, and can present antigens to cytotoxic T cells in vitro and in vivo. peptide. Analysis of post-mortem MS tissue revealed the presence of immunoproteasome expressing Sox-10^+^ cells in white matter lesions, suggesting that OPCs in human disease exhibit similar phenotypic changes. Therapeutic manipulation of this pathway could facilitate functional remyelination, as well as suppress OPC mediated inflammation, promote differentiation and reduce cytotoxic mediated cell death of OPCs.

## Results

### Adoptive transfer of effector T-cells inhibits remyelination by targeting OPCs

To better understand the mechanisms by which, IFNγ/IL-17 dual producing T-cells significantly inhibit remyelination, we adoptively transferred (AT) myelin reactive cytokine secreting T-cells into mice following 4 weeks of 0.2% CPZ, as previously described^36^. Detailed lineage tracing of OPCs was performed utilizing PDGFRα-CRE^ER^ x Rosa26-YFP mice (C57BL/6) (**supplementary fig1a**)^33^. In this way, the recombined population of OPCs mobilized to promote myelin repair following primary demyelination could be monitored through the entire process of remyelination and the influence of the subsequent effector T-cell transfer could be determined (**supplementary fig 1b-e**).

We analyzed YFP^+^-OPC differentiation and myelin content at 1 and 2 weeks after AT (**fig. 1a-d, supplementary fig. 2a,b**). In accordance with previous findings, we detected CD3^+^ cells at both time points in both CPZ and non-CPZ mouse corpus callosum following AT (**fig. 1a,b supplementary fig. 2a,b)**. Black Gold myelin staining revealed that remyelination was inhibited even at 2 weeks post AT. T-cells by themselves do not cause demyelination in non-CPZ corpora callosa (**fig. 1b, supplementary fig. 2b)**. A significant reduction in total YFP^+^ cells at both time points was found AT-CPZ (**fig. 1c,d)**. We analyzed the proportion of YFP^+^ oligodendrocyte lineage cells using the markers PDGFRα, and CC1. In AT-CPZ mice one week post adoptive transfer the two populations of OPC (YFP^+^/PDGFα^+^/CC1^−^) and intermediate oligodendrocytes (YFP^+^/PDGFRα^−^/CC1^−^) were significantly reduced in comparison to CPZ alone (**fig. 1a, c, supplementary fig. 2a**). The YFP+ mature oligodendrocyte population was significantly reduced in the AT-CPZ mice as compared to CPZ alone two weeks post adoptive transfer (**fig. 1 b, d, supplementary fig. 2b**). Since it would be expected that homeostatic OPC proliferation would maintain cell numbers, an explanation for our findings of reduced total numbers of YFP+ cells might be that the OPCs or oligodendrocyte lineage cells were undergoing cell death specifically in AT-CPZ mice, a hypothesis that we mechanistically pursued in subsequent experiments.

**Figure 1:**
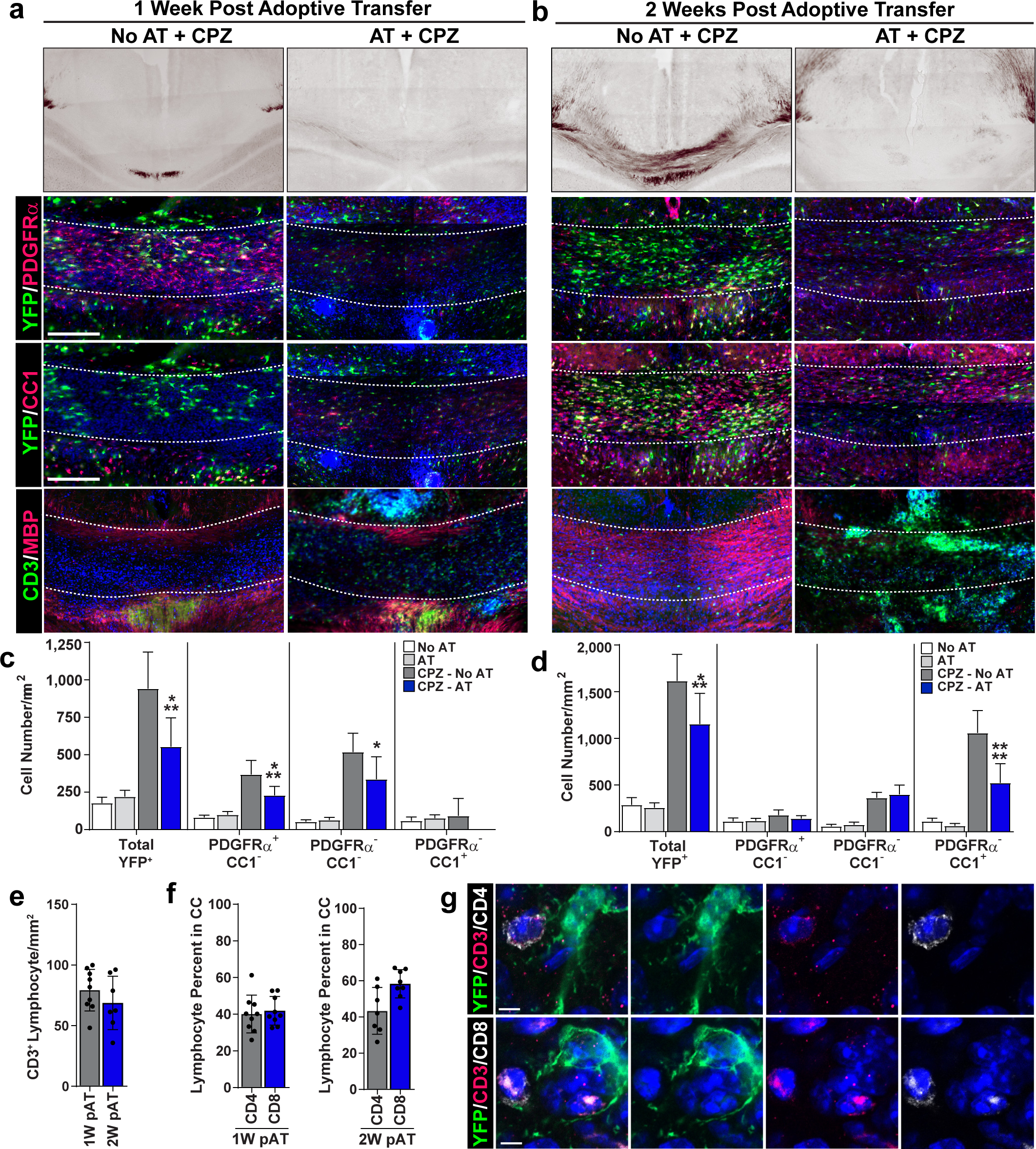
Adoptive transfer of effector T-cells inhibits remyelination by targeting OPCs. PDGFRα-Cre^ER^ x Rosa26-YFP bred to the C57BL/6 background were kept on a 0.2% CPZ diet for a total of 4 weeks. After 3 weeks CPZ, 4-hydroxytamoxifen (1 mg/mouse/day for 3 days) was injected to induce Cre recombination in PDGFRα expressing cells. Upon recombination PDGFRα expressing cells heritably express YFP regardless of differentiation status. Approximately 8-10 million MOG35-55 specific effector T-cells were isolated and purified from 2D2 TCR transgenic mice and were injected IP into recipient mice at 4 weeks. Simultaneously, the recipient mice were put back on a normal feed diet and were sacrificed 1-2 weeks after adoptive transfer. **(a)** 1-week (scale bar 400 μm) and **(b)** 2-week images of Black Gold myelin staining (top panel). Representative images of the corpus callosum (outlined by white dashed line) of brain sections. (**a, b 2^nd^ row**) stained with YFP (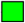) and PDGFRα (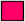) allowed tracking of recombined OPCs. Representative images of the corpus callosum of brain sections stained with YFP (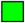) and CC1 (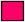) identified recombined mature oligodendrocytes (**a, b 3^rd^ row**). Representative images of the corpus callosum of brain sections stained with CD3 (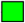) amd MBP(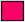) show the distribution of lymphocytes and myelin (4^th^ row). **(c-d)** Quantification of 1-week **(c)** and two-week **(d)** immunohistochemistry data to identify different stages of oligodendrocyte differentiation using the markers YFP, PDGFRα, and CC1. Oligodendrocyte lineage populations were compared between groups; No CPZ (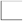; n=6, 8), No CPZ + AT (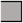; n=8, 8), CPZ (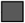; n=7, 16), and CPZ + AT (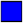; n=7, 11). Significance for quantified data **(c-d)** was assessed by one-way ANOVA analysis followed by Tukey’s multiple comparison analysis (α = 0.05, * ≤ 0.05, ** ≤ 0.01, *** ≤ 0.001, **** ≤ 0.0001). **(e)** Quantification of immunohistochemistry staining of total CD3 in the corpus callosum at 1-week and 2-week post adoptive transfer (**left**). **(f)** Of the total CD3 positive percent CD4 (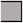; n=9, 7) and CD8 (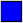; n=9, 8) was quantified at 1-week (**middle**) and 2-week (**right**). (g) Representative confocal images of CD3^+^/CD4^+^ T-cell interaction (**top**) and CD3^+^/CD8^+^ T-cell interaction (**bottom**) with YFP^+^ oligodendrocyte lineage cell.

In order to better understand the correlation between T-lymphocytes and remyelination we analyzed the number of CD3^+^ cells at one week and two weeks post AT and CPZ withdrawal (**fig. 1e**) and found that the density of T-cells in the corpus callosum were not significantly different between time points. While MBP staining was overall reduced in the AT-CPZ mice, the distribution of areas of demyelination and remyelination was variable throughout the corpus callosum of AT-CPZ mice, therefore we examined the relationship between CD3^+^ signal intensity and MBP rich or MBP sparse regions (**supplementary fig. 3a**). The signal intensities of MBP and CD3^+^ were negatively correlated (**supplementary fig. 3b**), supporting the hypothesis that CD3^+^ cells establish a microenvironment that is not conducive to remyelination. Taken together, these results suggest that IL-17/IFNγ producing T-cells disrupt remyelination independent of differentiation status of the OPC. CD4+ and CD8+ T-cell density in the corpus callosum were not significantly different at either the one week or two-week time point (**fig. 1f**) and both cells were found to interact with YFP+ oligodendrocyte lineage cells (**fig. 1g**).

### IFNγ and IL-17 inhibit OPC differentiation, but only IFNγ signaling promotes antigen cross-presentation and immunoproteasome pathways in OPCs

Since CD4^+^ T-cells exhibit their effector function primarily through cytokine production, we interrogated the influence of IFNγ and IL-17 on the process of OPC differentiation. Primary mouse OPCs were cultured *in vitro* following A2B5 positive selection (95-98% purity)^37^. OPCs were expanded with PDGF prior to cytokine treatment and the standard differentiation stimulus of T3 was initiated^38^. Differentiating OPC cultures were treated for a total of 96 hours prior to analysis of gene (**fig. 2a**) and protein (**fig. 2b**) expression. Both IL-17 and IFNγ significantly reduced MBP levels, while Sox-10 gene expression and Olig2 protein expression were not significantly altered. With the goal of ascertaining OPC functional outcomes beyond inhibition of differentiation, we hypothesized that IFNγ stimulation would alter the phenotype of OPCs, shifting from basal to inflammatory functions. Affymetrix microarray analysis of primary OPCs treated with T3 and IFNγ was conducted from 8-96 hours of treatment to capture gene expression changes throughout the entire process of OPC differentiation into mature oligodendrocytes, and reveal secondary gene expression changes (**fig. 2c,d**). IFNγ initiated a robust upregulation and downregulation of genes across multiple time points. Three main gene expression profiles were observed upon comparison with baseline PDGF-treated OPCs (**fig. 2d**). IFNγ delayed the influence of T3 on the upregulation or downregulation of gene expression, profiles 1 and 3, respectively. Profile 2 had a unique pattern, in which gene expression changes were directly linked to IFNγ treatment and independent of the pro-differentiation or pro-progenitor programs.

**Figure 2:**
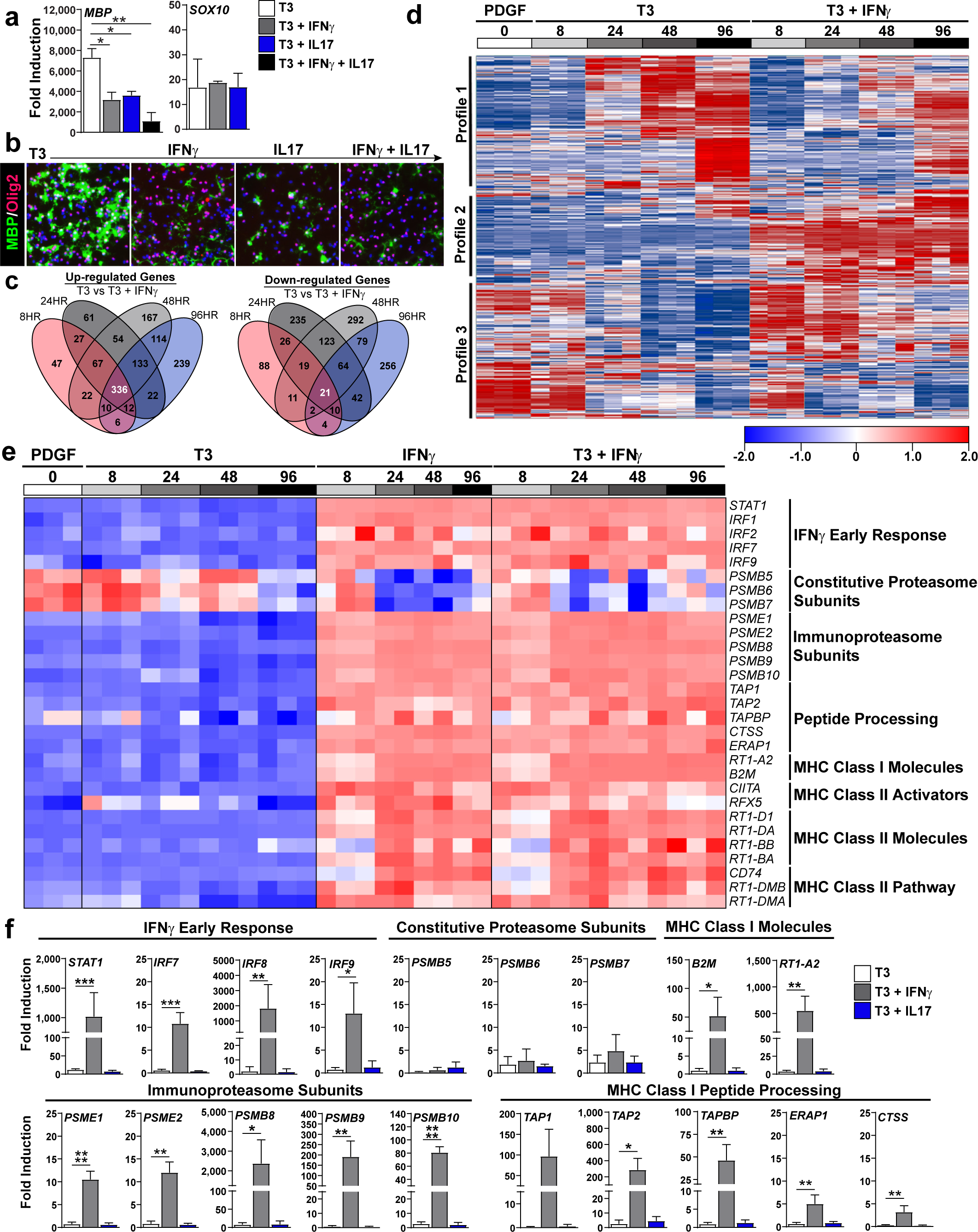
IFNγ and IL17 inhibit OPC differentiation genes while antigen cross-presentation and immunoproteasome pathways are only enriched under IFNγ stimulation. Isolated and PDGF (20 ng/mL) expanded postnatal rat OPCs (P4-P6) were differentiated with T3 (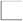), T3 + IFNγ (10 ng/mL; 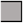), T3 + IL17 (50 ng/mL; 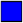), or T3 + IFNγ + IL-17 (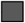) for 96 hrs prior to assessment using qPCR **(a)** and ICC staining for MBP and Olig2 **(b)**. Isolated and PDGF (20 ng/mL) expanded postnatal rat OPCs (P4-P6) were left undifferentiated (0 hrs; 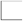) or differentiated with T3 (10 nM), IFNγ (10 ng/mL), T3 + IFNγ, for 8 hrs (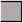), 24 hrs (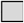), 48 hrs (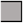) and 96 hrs (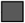) to compare gene expression between groups while obtaining information at multiple time points. **(c-e)** Affymetrix gene arrays analysis was performed from three independent biological replicates. **(c)** Venn diagram summarizing the number and overlap of up-regulated (top) and down-regulated (bottom) genes based on time point. **(d)** Global heat map of all probes clustered into three gene expression patterns comparing PDGF, T3, and T3 + IFNγ at all time points. **(e)** Targeted heat map comparing PDGF, T3, IFNγ, and T3 + IFNγ at all time points. Displayed genes were identified from GSEA analysis (supplement 3). **(f)** Quantitative PCR validation of OPCs differentiated with T3 (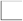), T3 + IFNγ (10 ng/mL; 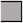) or T3 + IL17 (50 ng/mL; 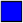) for 96 hrs prior to assessment. Error bars represent the standard deviation from 3 independent primary isolations and experiments. Significance for qPCR analysis was determined by one-way ANOVA analysis followed by Dunnett’s multiple comparison analysis where T3 + IFNγ or T3 + IL17 were compared to T3 alone control (P * ≤ 0.05, ** ≤ 0.01, *** ≤ 0.001, **** ≤ 0.0001).

To interrogate the signaling pathways altered in OPCs by IFNγ, we performed gene set enrichment analysis (GSEA; **supplementary fig. 4a,b**). Consistent with our differentiation data, oligodendrocyte markers were highly enriched in T3 treatment alone compared to T3 + IFNγ (**supplementary fig. 4a**). IFNγ treatment promoted the induction of genes that constitute antigen processing and cross-presentation signaling pathways at all time points analyzed (**fig. 2e and supplementary fig. 4b;** 24 hour and 48 hour data not shown). Of note, the immunoproteasome subunits that define the transition of the proteasome during antigen processing were significantly increased, while the constitutive proteasomal subunits were significantly decreased. Furthermore, both MHC class I and class II molecules were significantly augmented in response to IFNγ.

We performed quantitative PCR analysis to validate the microarray data and found that genes involved in antigen processing, MHC class I and MHC class II, were significantly increased in response to IFNγ (**fig. 2f, supplementary fig. 4 – gray bars**). However, the mechanism by which IL-17 orchestrates its inhibitory effect on OPCs is independent of the antigen processing and MHC class I/II signaling, since we were unable to detect upregulation of the genes that comprise these pathways in OPCs that had been exposed to IL-17 (**fig. 2f, supplementary fig. 5 – blue bars**). The results suggest that OPCs undergo dramatic phenotypic changes when exposed to IFNγ, adopting features commonly associated with immune cells, such as antigen processing and presentation.

### CNS restricted expression of IFNγ promotes CD8 infiltration and MHC class I and MHC class II expression in OPCs

To determine whether IFNγ promotes OPC antigen presentation *in vivo*, we first examined whether CNS specific expression of IFNγ would upregulate the gene expression profiles that we previously observed. GFAP/tTA transgenic mice were crossed with TRE/IFNγ transgenic mice^39,40^. The Tet-off system allows for controlled IFNγ expression from GFAP^+^ astrocytes upon removal of dietary doxycycline (DOXY OFF) without the need to induce CNS inflammation, and was previously shown to suppress endogenous remyelination the CPZ model^41,42^. Animals were provided 0.2% CPZ to induce demyelination and subsequently increase the pool of OPCs prior to corpus callosum micro-dissection and qPCR analysis of DOXY ON mice and DOXY OFF. Consistent with our *in* *vitro* gene expression data, we found that IFNγ induced upregulation of the genes involved in MHC class I and MHC class II antigen presentation (**fig. 3a**). To demonstrate the specific effect of IFNγ on OPCs, whole brain tissue was removed after two weeks of CPZ withdrawal and flow cytometry was used to investigate lymphocyte and OPC cellular populations under IFNγ and non IFNγ conditions *in vivo.* We compared OPC-H2Kb expression in DOXY ON versus DOXY OFF mice and found significant increases in OPC surface expression of H2Kb in mice induced to express IFNγ (**fig. 3b)**. Upon more detailed analysis of the DOXY OFF H2Kb expressing OPCs we found that a significantly higher proportion of H2Kb^+^ OPCs were also dual positive for pan MHC class II molecules (IA/IE) than in DOXY ON mice (**fig. 3c**). Next, we determined how CNS restricted IFNγ expression influenced T-cell and OPC percentages in the brain. While CD8^+^ T-cell percent was dramatically increased in response to IFNγ, the percent of CD4^+^ T-cell remained unchanged (**fig 3d**). We interrogated different populations of OPC based on the three markers PDGFRα, A2B5 and O4 with a parent gate of CD11b^−^/Olig2^+^. CNS expression of IFNγ significantly reduced total PDGFRα^+^, PDGFRα^+^/A2B5^+^, and PDGFα^+^/O4^+^ expressing OPCs (**fig. 3e**). Due to the observed influence on CD8^+^ T-cell percentages, their known cytotoxic effector function, the robust expression of MHC class I and the observed loss of OPCs we decided to focus on OPC-CD8^+^ T-cell interaction by mean of antigen presentation.

**Figure 3:**
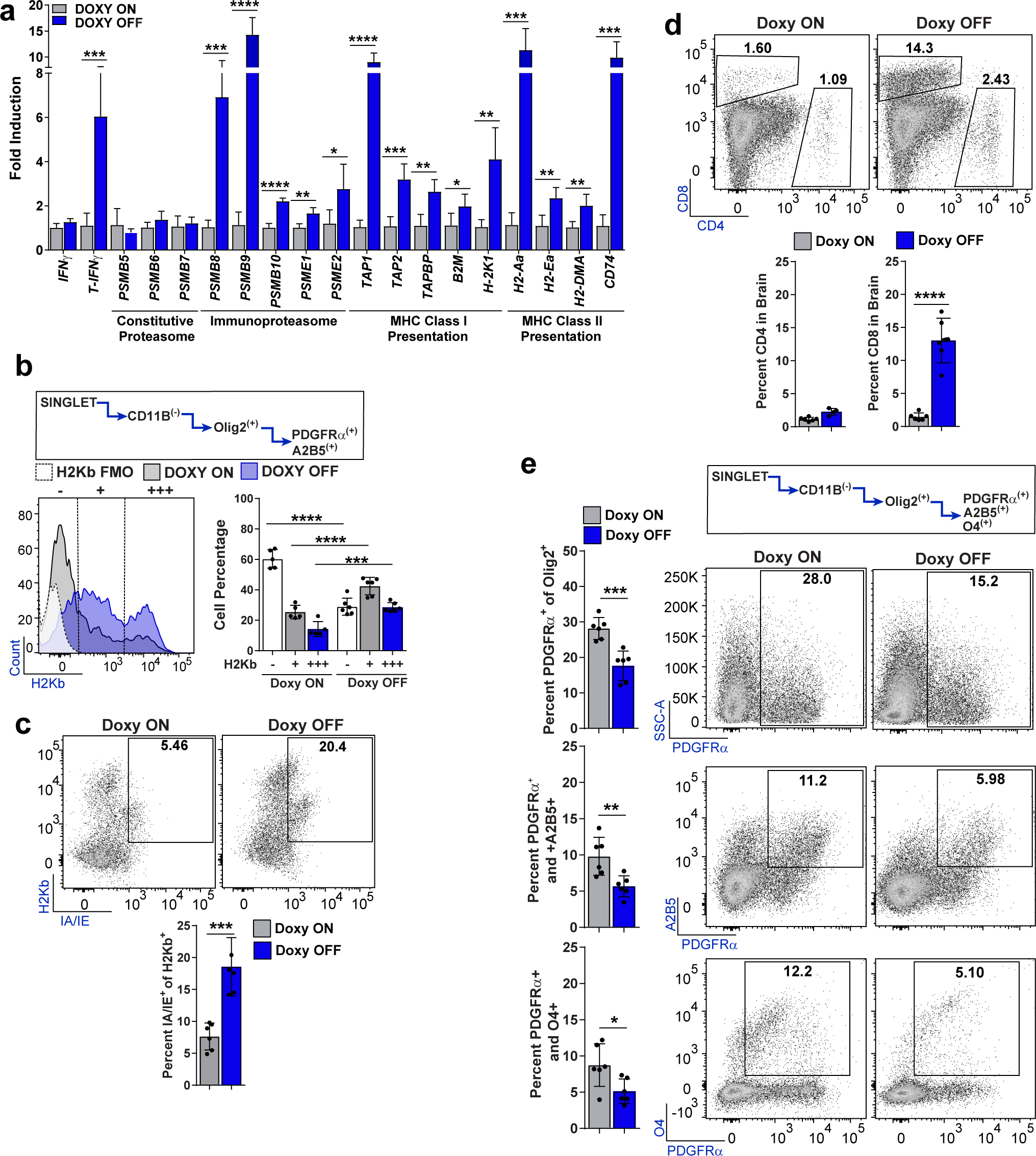
IFNγ signaling in the CNS causes CD8 recruitment and MHC Class I and II expression on OPCs. **(a)** TRE/IFNγ x GFAP/tTA mice were fed CPZ for a total of 6 weeks. At the start of CPZ mice were either kept on doxycycline (DOXY) (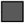; n=5) or removed from doxycycline (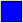; n=5) to induce astrocyte-derived IFNγ expression. Two weeks after replacement of normal diet mice were sacrificed and the corpus callosum was dissected and prepared for qPCR analysis. Significance for qPCR analysis was determined by two-tailed, unpaired t-test between doxycycline and no doxycycline conditions (P * ≤ 0.05, ** ≤ 0.01, *** ≤ 0.001, **** ≤ 0.0001). Error bars represent standard deviation **(b)** Whole brains from TRE/IFNγ x GFAP/tTA mice under the same experimental paradigm were isolated for flow cytometry analysis. The OPC population was determined by CD11b negativity and Olig2, A2B5, and PDGFRα positivity. H2Kb expression is shown in the histogram plot in which the staining control FMO (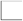; pooled) is compared to mice kept on doxycycline (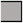; n=5) and mice removed from doxycycline (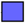; n=5). Significance for flow cytometry analysis was determined by two-tailed, unpaired Student’s t-test between doxycycline and no doxycycline conditions (P * ≤ 0.05, ** ≤ 0.01, *** ≤ 0.001, **** ≤ 0.0001). Error bars represent standard deviation. **(c)** The same gating strategy was used to determine MHC Class II (IA/IE) expression by H2Kb. Significance for flow cytometry analysis was determined by two-tailed, unpaired t-test between doxycycline (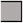; n=6) and no doxycycline (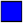; n=6) conditions were compared (P * ≤ 0.05, ** ≤ 0.01, *** ≤ 0.001, **** ≤ 0.0001). Error bars represent standard deviation. CD4^+^ and CD8^+^ populations were also analyzed within the whole brain tissue as determined by flow cytometry. The CD4+ population remains unchanged between doxycycline (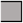; n=6) and doxycycline removal (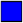; n=6). However, the removal of doxycycline significantly increased the number of CD8^+^ cells in the brain. Significance for flow cytometry analysis was determined by twotailed, unpaired Student’s t-test between doxycycline and no doxycycline conditions were compared (P * ≤ 0.05, ** ≤ 0.01, *** ≤ 0.001, **** ≤ 0.0001). Error bars represent standard deviation. (**e**) Percent OPC numbers were also analyzed for PDGFRα, A2B5 and O4. PDGFRα^+^ (**top**), PDGFRα^+^/A2B5^+^ (**middle**) and PDGFRα^+^/O4^+^ (**bottom**) OPC percentages were all significantly decreased in animals in which doxycycline was removed. Significance for flow cytometry analysis was determined by two-tailed, unpaired Student’s t-test between doxycycline and no doxycycline conditions were compared (P * ≤ 0.05, ** ≤ 0.01, *** ≤ 0.001, **** ≤ 0.0001). Error bars represent standard deviation.

### IFNγ stimulation induces peptide processing and presentation in OPCs *in vitro*

To address the functionality of the immunoproteasome and antigen presentation signaling pathways in OPCs, we examined MHC class I (H2Kb in C57/BL6 mice) expression and presentation of exogenous MHC class I restricted ovalbumin (OVA) peptide (OVA257-264) following IFNγ stimulation. We treated primary mouse OPCs with IFNγ for 12 hours and then provided OVA257-264 for 8 hours to allow for engulfment and peptide binding in the MHC class I groove, followed by presentation on the cellular surface (**fig. 4a**). To identify presentation of OVA257-264 peptide on the MHC class I molecule, we utilized an antibody that has specific affinity for OVA257-264 loaded H2Kb molecules (H2Kb-OVA)^43^. We observed high expression of H2Kb-OVA in PDGFRα immunoreactive OPCs following IFNγ stimulation, but not with a control MHC class I restricted peptide (MOG37-50) or in no cytokine conditions.

**Figure 4:**
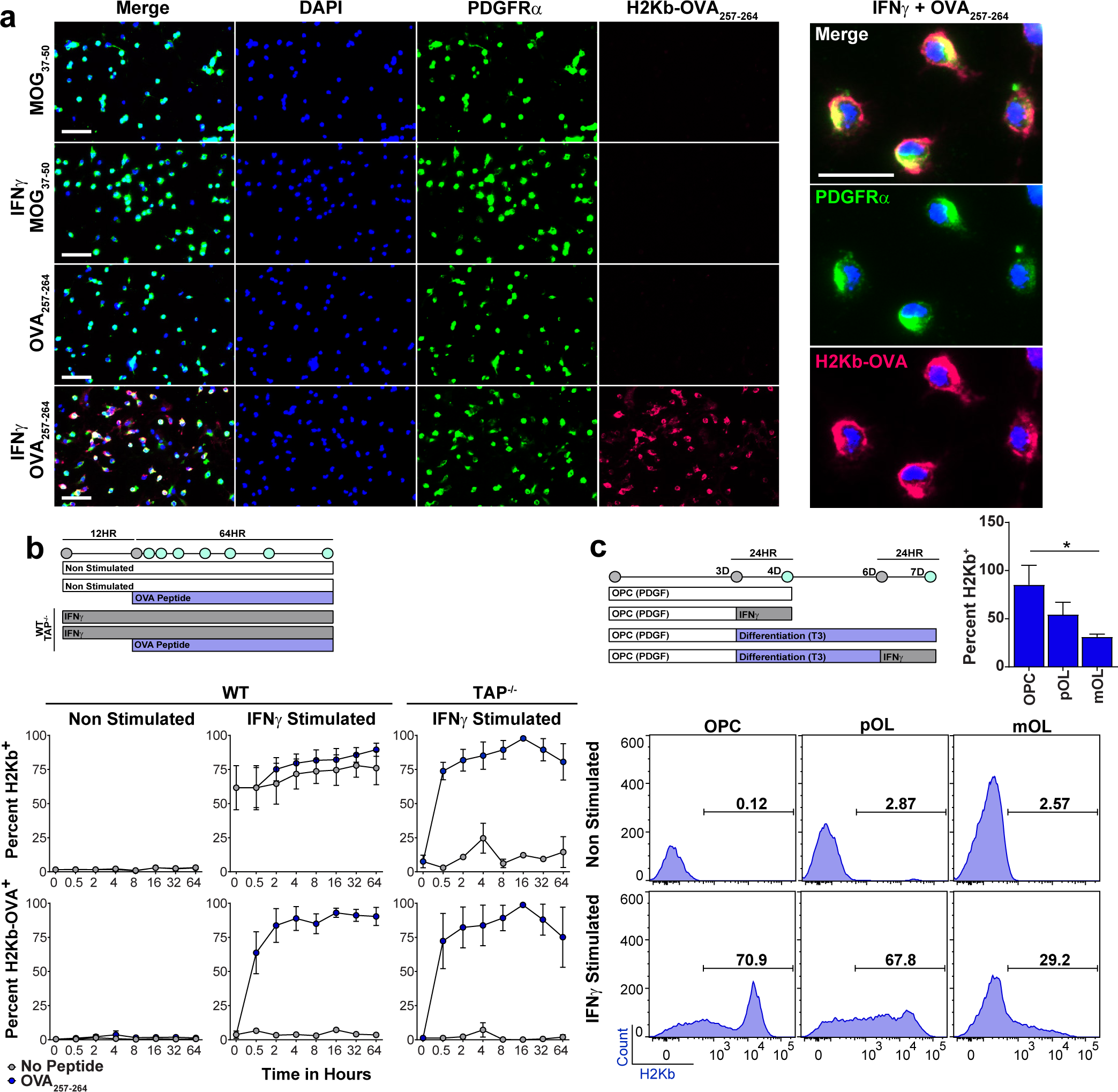
IFNγ stimulation induces stable and rapid peptide processing in OPCs both *in vitro* and *in vivo*. **(a)** Immunocytochemical staining of primary C57BL/6 mouse OPCs, derived from OT1 TCR transgenic mice, which were cultured under PDGF conditions or with IFNγ (10 ng/mL) for 12 hrs prior to addition of the MHC class I restricted MOG37-50 peptide (50 μg/mL) or OVA257-264 peptide (50 μg/mL). Ovalbumin class I peptide presentation on OPCs was determined using PDGFRα (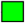) and H2Kb-OVA257-264 (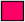) antibodies. The right panel shows high magnification (100 μm) of IFNγ + OVA257-264 culture. **(b)** Time course experiments of WT and TAP^−/−^ OPCs cultured with/without IFNγ and with/without OVA257-264 peptide. Treatment of IFNγ was completed for 12 hours prior to the addition of no peptide (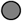) or OVA257-264 peptide **(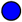(c)** MHC class I expression was determined from three stages of oligodendrocyte lineage cells either unstimulated or stimulated with IFNγ for 24 hrs. Oligodendrocyte lineage populations were defined by PDGFRα, A2B5, and O4 markers under proliferative or differentiating conditions.

MHC class I restricted peptides can be processed and presented through either the cytosolic pathway that is dependent on the transporter associated with antigen processing (TAP1) taking antigen processed by the immunoproteasome and transferring it to the ER, or a vacuolar pathway that is independent of immunoproteasome and TAP1 transport^44–50^. In order to further explore the dynamics of antigen processing by OPCs, we next determined the processing/presentation speed and stability of the OVA257-264 peptide on MHC class I molecules. After addition of OVA257-264 peptide to OPC cultures, we examined the percentage of OPCs presenting H2Kb-OVA (**fig. 4b**) and quantified the mean fluorescence intensity detected (**supplementary fig. 6b**) in WT and TAP1^−/−^ mice. Within 30 minutes of adding OVA257-264 to the cultures, over 50% of the IFNγ stimulated OPCs presented the peptide. While the percentage of OPCs expressing H2Kb-OVA remained stable at 80-90% between 2-64 hours, the mean fluorescence intensity continued to increase through 32 hrs. The dynamics of OPC peptide presentation were comparable to bone marrow derived dendritic cells (data not shown)^51^. One caveat of this experiment is that OVA257-264 peptide does not require cytosolic MHC class I processing, as shown by the ability of TAP1^−/−^ OPCs to express surface H2Kb-OVA (**fig 4b, right, supplementary fig. 6a)**. However, given that MHC class I is only expressed on the surface after intracellular peptide loading, it is unlikely that OVA peptide was externally coating MHC class I molecules, because H2Kb is detected on the surface of TAP1^−/−^ OPCs only after OVA257-264 is provided. Therefore, these results suggest that OPCs engulfed the peptide and internally loaded it onto MHC class I molecules through a TAP1^−/−^ independent mechanism termed the vacuolar pathway, as has been described^50^. We subsequently verified the distinct processing of whole OVA protein as being dependent on the cytosolic pathway, in functional experiments detailed below (**fig. 5d**). Since oligodendrocytes have previously been shown to present antigen, we compared the capacity of IFNγ stimulated OPCs, intermediate oligodendrocytes, and mature oligodendrocytes to present MHC class I^52^. OPCs were 97% MHC class I^hi^ after 24 hours of IFNγ stimulation compared to intermediate (4%) and mature oligodendrocytes (2%) (**fig. 3c**). The OPCs that express MHC class I under IFNγ stimulation also express MHC class II (**supplementary fig. 7**).

**Figure 5:**
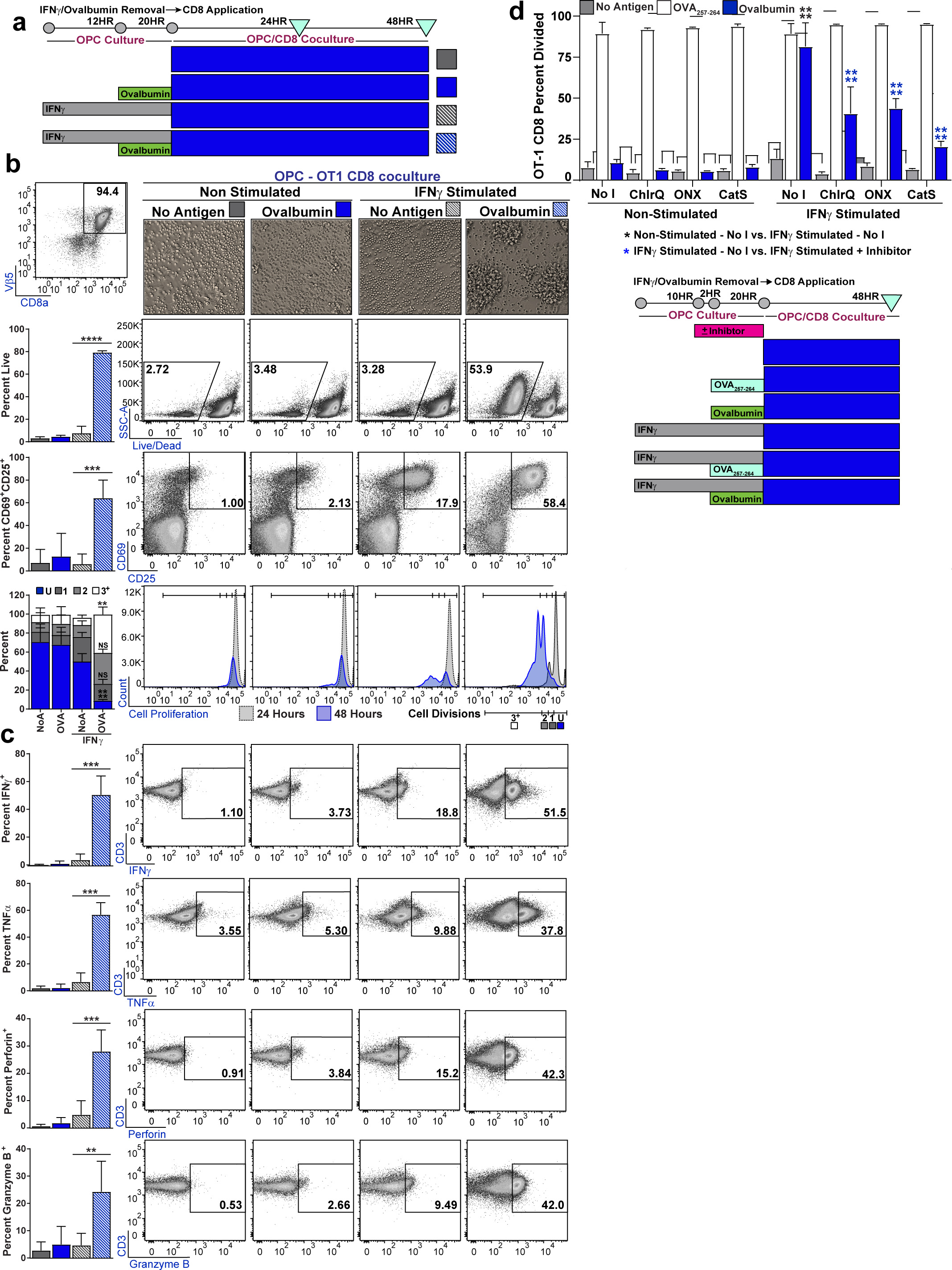
OPCs cross present ovalbumin protein and activate OVA-specific CD8^+^ T-cells. **(a)** Timeline and experimental design. OPCs were cultured in PDGF to inhibit differentiation and stimulated with IFNγ (10 ng/mL) for 12 hours prior to Ovalbumin (500 μg/mL) protein addition. Both IFNγ and Ovalbumin protein were incubated with OPCs for a total of 8 hours prior to washing the cultures to remove unprocessed Ovalbumin. OT-1 CD8^+^ T-cells were isolated by magnetic sorting then stained with Cell Proliferation Dye eFluor 450 (10 μM) prior to initiation of CD8/OPC co-culture. 24-48 hours after the start of the co-culture CD8 were analyzed for activation. **(b top left)** CD8 percentage, at 24 hours after the coculture was initiated, was determined using Vβ5, CD3, and CD8, and subsequently used as a parent gate for all flow plots in the figure. **(b)** CD8 morphology (**top**), survival (**2^nd^ row**), activation status using CD25^+^/CD69^+^ (**3^rd^ row**), and proliferation (**bottom**) with quantification (left to each panel) of OT-1s cultured with OPCs; no peptide (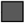), Ovalbumin (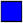), IFNγ + no peptide(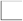), and IFNγ + Ovalbumin (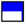). **(c)** Timeline of experimental methodology and treatment times (**bottom**), No inhibitor (No I) or each inhibitor were added to OPC culture for two hours prior to the addition of OVA257-264 or Ovalbumin, and then left in wells during processing and presentation phase. Chloroquine, an endosomal maturation inhibitor was added at 100 μM. ONX-0914, β5i/PSMB8 inhibitor was added at 30 nM. Cathepsin S (protease involved in antigen processing) inhibitor was added at 10 nM. **(top)** 48 hrs after CD8 co-culture was initiated cell proliferation of the Vβ5, CD3, CD8a population of CD8s was analyzed for cell division and total percent divided. Significance for all quantified data was either assessed by two-way ANOVA analysis followed by a Tukey’s multiple comparison analysis (P * ≤ 0.05, ** ≤ 0.01, *** ≤ 0.001, **** ≤ 0.0001). Error bars in all graphs represent standard deviation from 3 experiments. **(d)** Cytokine and granular protein profiling of OT-1s; IFNγ (**top**), TNFα (**2^nd^ row**), perforin (**3^rd^ row**) and granzyme B (**bottom**). Significance for all quantified data was assessed by one-way ANOVA analysis followed by Tukey’s multiple comparison analysis (P * ≤ 0.05, ** ≤ 0.01, *** ≤ 0.001, **** ≤ 0.0001). Error bars represent standard deviation for 5 biological replicates.

### OPCs cross-present ovalbumin protein and activate OVA-specific CD8^+^ T-cells

To further explore the functionality of OPC antigen presentation, we developed an OPC-CD8 co-culture system. OPCs were treated with IFNγ for 12 hours prior to OVA protein administration. Use of the full-length OVA protein necessitates engulfment, immunoproteasome processing and TAP1/2 dependent ER transport to successfully achieve surface expression of OVA peptide on MHC class I. OT-1, OVA257-264 peptide MHC class I restricted TCR transgenic mice were utilized to examine CD8 activation^53,54^. (**fig. 5a**). At 24 and 48 hours after co-culture initiation, CD8^+^ cell (selected by CD8a, CD3 and Vβ5; **fig. 5b, top left**) activation was examined by morphology, survival, CD25/CD69 immunoreactivity and proliferation (**fig. 5b**). Consistent with an activated phenotype, only OT-1 CD8s co-cultured with IFNγ stimulated/ovalbumin OPCs appeared clustered under phase contrast and had significantly higher survival, a higher percentage of CD25^+^/CD69^+^ cells and significantly more cell proliferation at 48 hours compared to controls. Importantly, OT-1 CD8s co-cultured with IFNγ/ovalbumin OPCs were similar in their ability to survive and proliferate when compared to splenocytes presenting processed OVA (**supplementary. fig. 8a**).

Complete CD8 cytotoxic T-cell (CTL) activation results in cytokine and granular protein, production providing CD8s with their effector function^18–20^. We investigated the ability of IFNγ-ovalbumin OPCs to induce OT-1 CTLs and found that significantly more CD8s were induced to produce IFNγ, TNFα, perforin, and granzyme B when compared to the other experimental conditions (**fig. 5c)**. The OPC induced CD8 cytokine production was comparable to the splenocyte (containing classical antigen presenting cells) induced CD8 cytokine production of IFNγ and TNFα (**supplementary fig. 8b**). These experiments indicate that IFNγ stimulated OPCs engage in antigen cross-presentation through the engulfment, processing, and surface presentation of antigen, enabling OPCs to activate CD8^+^ CTLs.

Furthermore, pharmacological blockade of different stages of the cytosolic MHC class I pathway using; Chloroquine (endosomal processing inhibitor), ONX-0914 (immunoproteasome β5i subunit inhibitor), or Cathepsin S (protease involved in antigen processing) inhibitor had no effect on OPC-OVA257-264 peptide presentation (vacuolar pathway), as determined by the OT-1 CD8 proliferation assay, while all three significantly hindered the ability of OPCs to engulf, process and present OVA full-length protein (cytosolic pathway). These results are consistent with the hypothesis that peptides can processed through the above described alternative vacuolar pathway, but whole protein requires the classical cytosolic processing machinery (**fig. 5d**).

### Caspase3/7 activity significantly increases in OPCs that activate OT-1 CD8

One of the consequences of CD8 activation is cytotoxic targeted cell death. This could be especially important in the context of demyelination in inflammatory diseases such as MS, as a potential mechanism for remyelination failure if OPCs were aberrantly targeted by CTLs. In addition to protease-mediated cytotoxicity, contact-dependent Fas-Fas Ligand surface interaction allows CD8s to induce apoptosis of its target while minimizing off-target effects^21–23^. Although perforin/granzyme and Fas/FasL intracellular signaling can be divergent, both pathways have been shown to promote apoptosis through caspase 3/7 activation^55–58^. To determine whether OPC-activated CD8s were in turn cytotoxic, we first examined if OPCs exhibited surface expression of Fas by flow cytometry (**fig. 6a**). A significantly higher percentage of OPCs expressed surface Fas after IFNγ treatment compared to non-stimulated OPCs. We used live cell imaging of OPC-CD8 co-cultures to measure induced caspase3/7 activity of OPCs. OT-1 CD8s cultured in isolation exhibited very little caspase 3/7 activity (**fig. 6b,d**). OPCs either non-stimulated or stimulated with IFNγ at baseline, 24 hours, and 48 hours had similar levels of caspase 3/7 as active OPCs (**fig. 6c,d**; white mask), whereas the IFNγ/ovalbumin conditioned OPCs that robustly activated OT-1 CD8s had significantly more cell death when compared to IFNγ/no antigen OPCs (**fig. 6c,d**). We analyzed the caspase 3/7 active population by flow cytometry and found greater H2Kb and Fas expression in this population compared to IFNγ stimulated/no antigen (**fig. 6e**). These experiments confirmed reciprocal effects of the cytotoxic CD8s on OPCs and identified a potential mechanistic pathway leading to cell death that could account, in part, for the observed OPC depletion in vivo.

**Figure 6:**
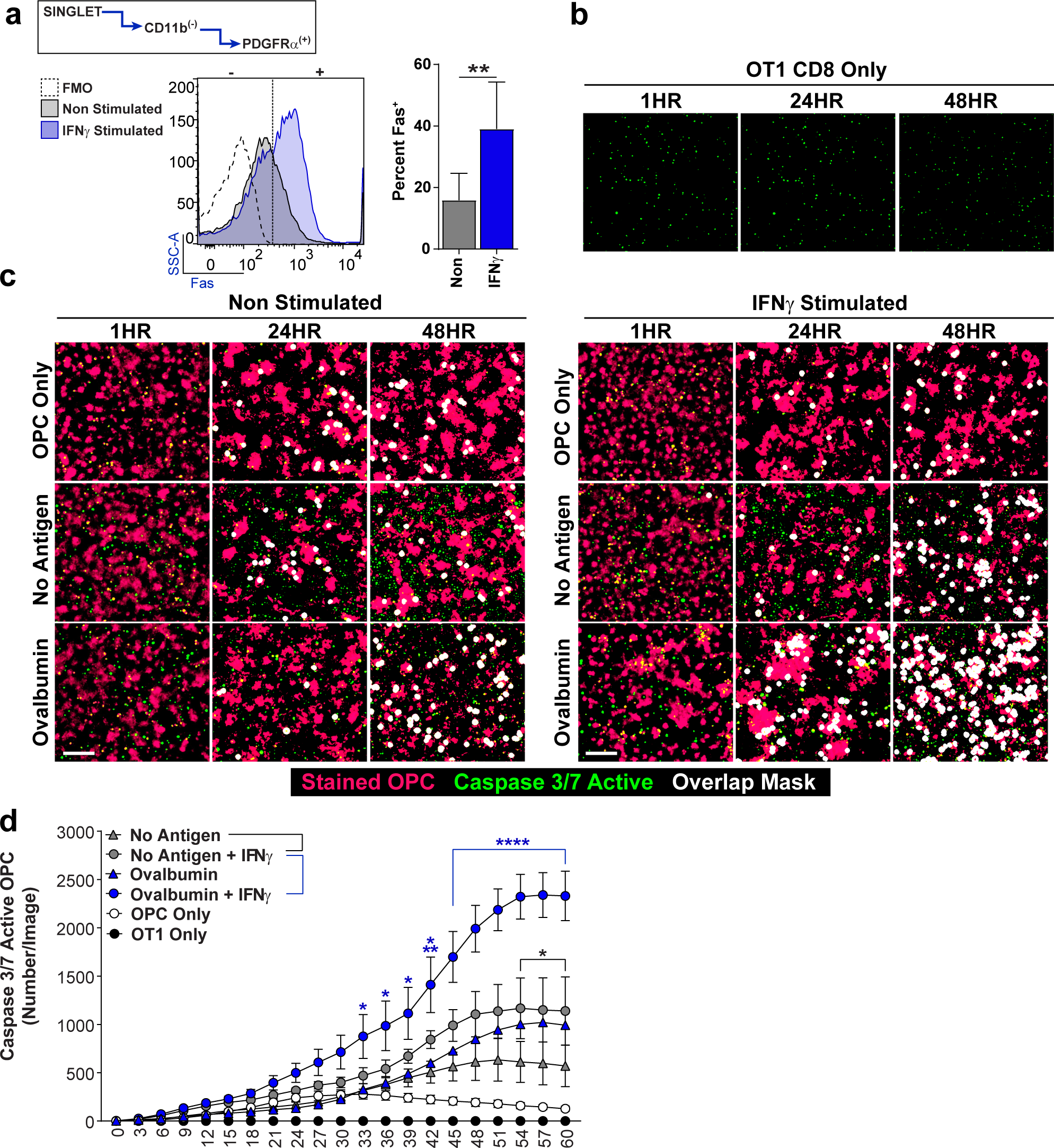
Caspase3/7 activity significantly increases in OPCs that activate OT-1 CD8. **(a)** Fas surface expression on OPCs, either not stimulated (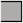) or stimulated with IFNγ (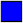), as determined by flow cytometry analysis for PDGFRα^+^/CD11b^−^ (FMO; 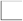). Significance was determined by unpaired, two-tailed Student’s t-test (P = 0.0046). Error bars represent standard deviation from 5 independent experiments. **(b)** OT-1 CD8s isolated and cultured for a total of 60 hours in the presence of activated caspase 3/7 detection reagent (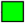). Representative images for 0 hrs, 24 hrs, and 48hrs are shown in the figure and quantified up to 60 hours (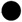) **(d).** **(c)** Time-lapse imaging of OPC-CD8 co-cultures in which OPCs were cultured under the following conditions prior to CD8 addition; no peptide (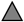), Ovalbumin protein (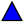), IFNγ stimulated OPC only (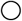). IFNγ + no peptide (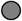) and IFNγ + Ovalbumin protein (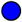). OPCs were stained (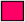) and caspase3/7 reagent (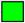) was added to identify OPCs that became caspase active (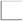). **(d)** Quantification of time-lapse images from 0 – 60 hrs. Significance was determined by 2way ANOVA followed by Tukey’s multiple comparisons test (P * ≤ 0.05, ** ≤ 0.01, *** ≤ 0.001, **** ≤ 0.0001). Error bars represent standard deviation from 3 experiments.

### OPCs express MHC class I loaded with myelin peptide and are targeted to die *in vivo*

To examine the consequences of MHC class I presentation *in vivo* we utilized two different strains of mice, each with unique advantages. First, we adoptively transferred 2D2^59^ MOG35-55-specific T-cells into the *PDGFRα-CRE^ER^ x Rosa26-YFP* cuprizone fed mice, which enabled us to trace OPCs with a reporter and have greater certainty of OPC-specific MHC class I expression CD8 CTL mediated cytotoxicity (**supplementary fig. 9a**). This model also has a detectable CD8 infiltrate even though it is induced with CD4s^35^. As previously described, we found significantly higher EAE scores in AT-CPZ-mice compared to no-AT-CPZ (**supplementary fig. 9b**). In addition, these mice showed pronounced and significantly more CD3^+^ lymphocytes (**supplementary fig. 9c**). H2Kb and Fas expression within the total OPC population was significantly higher in the mice with AT versus no-AT (**supplementary fig. 10a).** We gated on the YFP^+^/PDGFRα^+^/A2B5^+^ population in AT-CPZ and no AT-CPZ control and found a significant proportion of both YFP^+^ and YFP^−^ OPCs that were caspase 3/7 active (**fig. 7a**). Of note there were significantly fewer YFP positive OPCs in mice that received the AT as determined by flow cytometry making this result consistent with previously shown immunohistochemistry analysis (**fig. 7a**). We extended the analysis of this population and interrogated cell death, H2Kb expression, and Fas expression. The majority of the YFP^+^/caspase 3/7 active population were labeled dead, H2Kb positive and Fas positive (**supplementary fig. 10b**).

**Figure 7:**
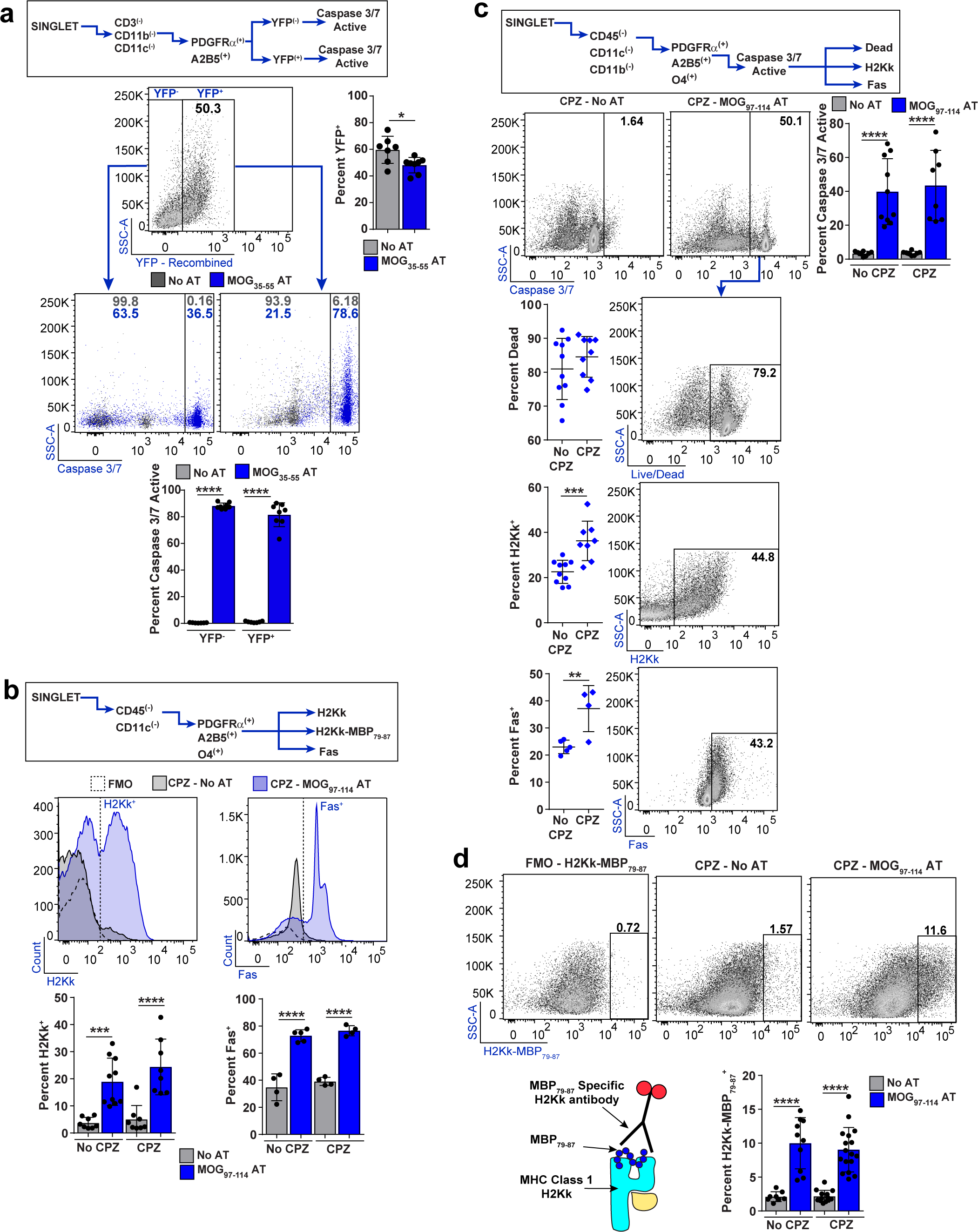
OPCs express MHC class I, MHC class I loaded with myelin peptide, and are targeted to die *in vivo* in two different animal strains. C57BL/6 PDGFRα-Cre^ER^ x Rosa26-YFP were kept on a CPZ diet for a total of 4 weeks. After 3 weeks, 4-HT (1 mg/mouse/day) was injected to induce Cre recombination in PDGFRα expressing cells and MOG35-55 reactive T cells were adoptively transferred (AT). Cells were isolated ex vivo and analyzed by flow cytometry. **(a)** The OPC population was distinguished based on YFP^−^ and YFP^+^ expression and analyzed for caspase 3/7 activity, quantification below; CPZ (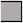) and CPZ + MOG35-55 AT (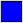). **(b-d)** Alternatively, C3HeB/FeJ donor mice were immunized with whole recombinant rat MOG1-125 to induce EAE, and myelin-specific CD4 T-cells were isolated *ex vivo* and reactivated with peptide. Syngeneic recipient mice were fed a CPZ diet for 6 weeks before cultured myelin reactive CD4 MOG97-114 specific cells were adoptively transferred. **(b)** H2Kk (**left**), and FAS (**right**) expression from the OPC population was determined by flow cytometry analysis with quantification below each flow histogram; no AT (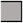; n = 8, 8, 4), MOG97-114 AT (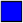; n = 9, 10, 5), CPZ + no AT (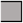; n = 8, 12, 4) and CPZ + MOG97-114 AT (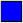; n = 8, 17, 4). Statistical significance was determined by one-way ANOVA analysis followed by Tukey’s multiple comparisons. Error bars represent the standard deviation. **(c)** Caspase 3/7 activity was determined by flow cytometry. Further population analysis was done from the caspase 3/7 active population by Live/Dead staining (**top**), H2Kk expression (**middle**) and FAS expression (**bottom**). **(d)** H2Kk-MBP79-87 staining from OPCs with FMO staining control and quantification below. Significance for all quantified data was either assessed by one-way ANOVA analysis followed by Tukey’s multiple comparison analysis or unpaired, two-tailed t-test (P* ≤ 0.05, ** ≤ 0.01, *** ≤ 0.001, **** ≤ 0.0001). Error bars in all graphs represent standard deviation.

We utilized an additional adoptive transfer model to further validate OPC antigen presentation and to measure MBP79-87 peptide in the MHC class I groove using an antibody specific for this peptide on H2Kk in C3HeB/FeJ mice^52,60^. This system was previously employed to show that TNF/iNOS producing dendritic cells (Tip-DCs) and mature oligodendrocytes mediate determinant spreading in EAE (**supplementary fig. 11a-g**)^52^. Examination of H2Kk and Fas expression were consistent with the previous model in which the AT with and without CPZ fostered increased expression of both markers in OPCs (PDGFRα^+^/A2B5^+^/CD11b^−^/CD45^−^) (**fig. 7b**). MOG97-114 AT caused a significantly higher proportion of OPCs with induced caspase 3/7 activity (**fig. 7c**). When comparing AT-caspase 3/7 active OPCs with CPZ-AT-caspase3/7 active OPCs we found there was no difference in the percentage of dead cells, but there was a significantly higher proportion of H2Kk and Fas immunoreactive cells in mice fed CPZ (**fig. 7c**). One possible explanation for this difference could be the level of IFNγ production by lymphocytes between AT alone and AT-CPZ. In accordance, we previously found that CPZ shifts the lymphocyte brain infiltrate from higher percentages of IL-17 producing CD4s to predominantly dual producing IFNγ/IL-17 CD4s^35^. Notably, we were able to detect MBP79-87 MHC class I restricted peptide expression within OPCs in animals that had CNS inflammation generated following AT (**fig. 7d**). The detection of the OPC MBP peptide presentation suggests that under inflammatory conditions, OPCs can mediate determinant spreading. These data show that in an inflammatory setting, OPCs can be induced to function as antigen presenting cells *in vivo* and present myelin peptide on MHC class I molecules. Furthermore, at peak disease, a significant percentage of OPCs are targeted to undergo caspase 3/7 mediated death.

### MS white matter lesions have significantly higher PSMB8 expression in oligodendrocyte lineage cells

To understand the clinical relevance of these observations and expand our studies of OPC phenotypic changes to human tissue samples, we conducted immune-histochemical (IHC) staining of human brain tissues collected from control and MS patients through autopsy. Using myelin proteolipid protein (PLP) staining, regions of normal white matter (NAWM) and demyelinated white matter (WML) were identified. Staining for PSMB8, a subunit specific to the immunoproteasome, was used to define cells that had activation of the MHC class I pathway (**fig. 8a**). Highly intense and prevalent staining for PSMB8 was observed in WML areas compared to both control and NAWM tissues. Furthermore, upon quantification, there were significantly more cells that were PSMB8^+^ in the WML regions compared to NAWM and healthy control brain white matter (**fig. 8b**). To ascertain which cells within the lesion were expressing PSMB8, Sox-10, an oligodendrocyte lineage marker was co-immuno stained in parallel sections (**fig. 8c**). There were significantly more PSMB8^+^/Sox-10^+^ cells in WML compared to control and NAWM (**fig. 8d**), indicating that oligodendrocyte lineage cells within human white matter MS lesions upregulate a key component of the immunoproteosome pathway. Taken together, the results provide evidence that oligodendrocyte lineage cells in the MS white matter lesions upregulate the immunoproteosome pathway specifically in areas of failed remyeliantion.

**Figure 8:**
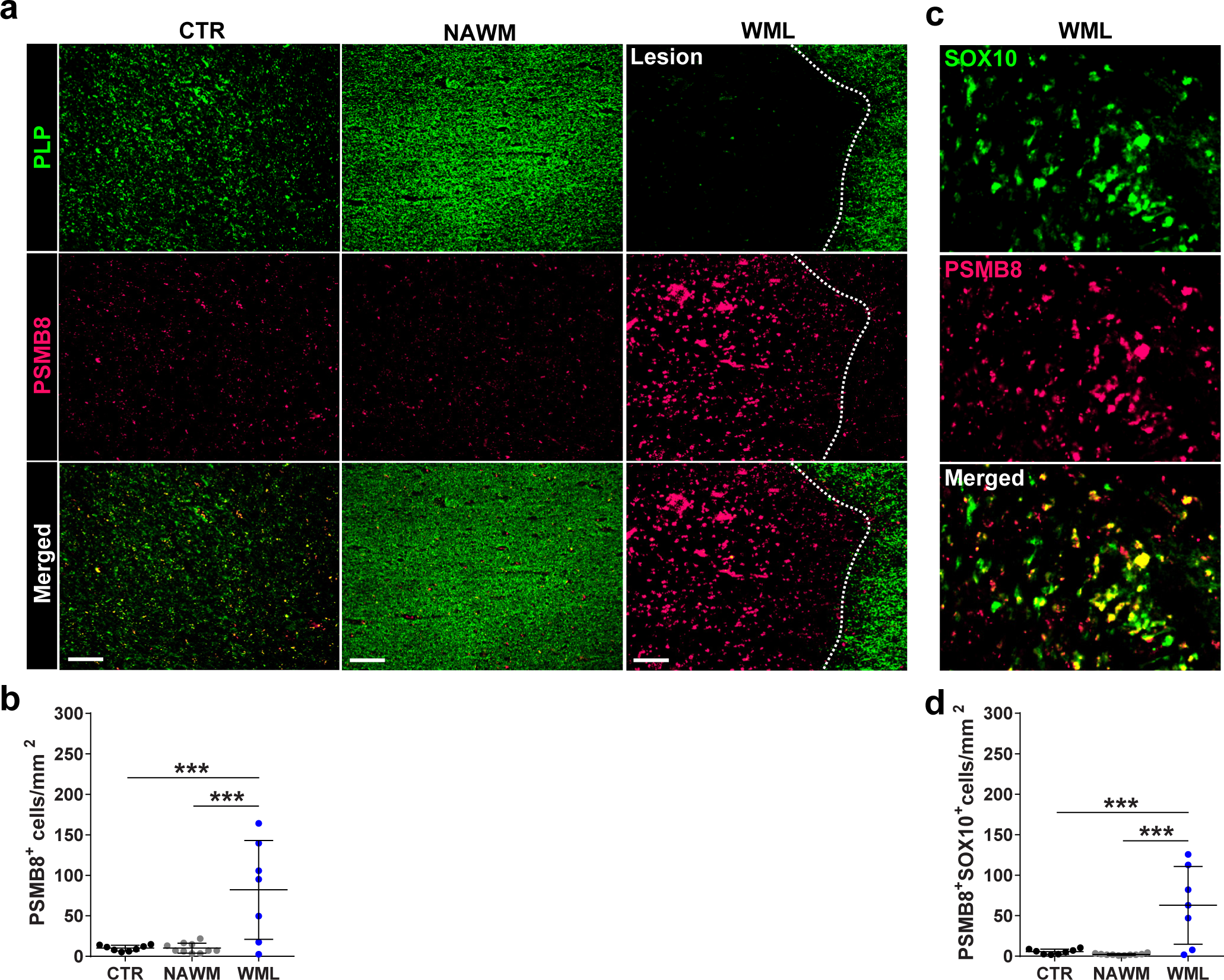
MS white matter lesions have significantly higher PSMB8 expression in oligodendrocyte lineage cells. **(a)** Postmortem tissue from healthy controls or MS patients (NAWM and WML) was analyzed for the immunoproteasome specific subunit, PSMB8 (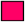) and PLP (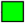) (scale bar 200 μm). **(b)** PSMB8 staining (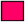) colocalization with SOX10 staining (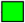). **(c-d)** Quantification of imaging. A total of 8 healthy controls, 10 NAWM and 7 WML sections were counted. Significance was determined by one-way ANOVA analysis followed by Tukey’s multiple comparison analysis (P * ≤ 0.05, ** ≤ 0.01, *** ≤ 0.001, **** ≤ 0.0001). Error bars in all graphs represent standard deviation.

## Discussion

Our findings demonstrate that OPCs can function as antigen presenting cells in the inflamed CNS, thereby not only failing to facilitate remyelination, but actually propagating continued inflammation. Specifically, we show that IFNγ induces expression of the MHC class I antigen presentation pathway in OPCs conferring an ability to activate CD8^+^ CTLs, which can in turn kill the OPCs as target cells. Our data also show that OPCs cross present exogenous proteins on class I molecules, a process that has previously been attributed to subsets of professional antigen presenting cells such as dendritic cells^49,61^. We demonstrate that OPCs switch from the constitutive proteasome to the immunoproteasome when exposed to IFNγ. This process can be suppressed by pharmacological inhibitors of the cytosolic processing machinery associated with this pathway. Finally, the marked upregulation of the critical immunoproteasome subunit PSMB8 on Sox-10 lineage cells only in demyelinated plaques of MS white matter lesions, suggests this process may be a critical component of the chronically demyelinated lesions of people with longstanding MS.

Previous studies have suggested that OPCs are responsive to a variety of inflammatory cues, which can be either pro-inflammatory, disease causing cytokines or, alternatively, immune regulatory and tissue healing. Two recent studies highlighted the regenerative role of M2 macrophages and Tregs in facilitating OPC differentiation and endogenous remyelination^7,9,62^. However, Tregs are known to be deficient in MS, and the disease is thought to be driven by Th1/Th17 biased immune cells^63–67^. Therefore, our focus has been on how effector T-cells might cause damage or hinder repair. Several reports have documented that OPCs can be suppressed or killed by TNFα^68,69^, IFNγ^25–30,70^ and IL-17^5,6^. IL-17 signaling through Act-1 has been shown to be critical in the pathogenesis of EAE specifically in OPCs^5,6,8^. Gene expression changes resulting from IL-17 signaling interfere with OPC maturation and promote production of inflammatory mediators including chemokines facilitating the recruitment of immune cells^6,8^. Our report is unique in that we show that the antigen cross presentation pathway in OPCs is triggered by IFNγ signaling, which was not previously envisioned as a mechanism of remyelination failure. We found that this IFNγ controlled pathway was not induced by IL-17. These findings suggest specificity of IFNγ for control of immunoproteasome/antigen presentation; however, there could be other cytokines that control these processes in OPCs. In support of this, A1 astroglia were recently shown to upregulate the immunoproteasomal subunit, PSMB8, following stimulation with IL-1α, TNFα and C1q^71^, raising the possibility that neuroinflammation stimulates a coordinated response among different microglia.

Neuroinflammation by T lymphocytes occurs in a two-step process^72–76^. The first step of peripheral priming of lymphocytes by dendritic cells ensues predominantly in secondary lymphoid tissues, after which T-cells home to the CNS. The second step happens soon after lymphocyte infiltration, in which T-cell are reactivated in either the subarachnoid or perivascular space, depending on the point of entry. Professional APCs, both dendritic cells, and macrophages residing in these locations re-activate the T-cells that subsequently enter into the parenchyma. The mechanisms that enable sustained T-cell activation within the parenchyma are not well understood, in particular, the duration of residence and the requirements for continued activation and survival. While knock out of MHC class II from CNS resident cells reduces infiltration of CD4^+^ T-cells and ablation of IL-17 signaling in OPCs was shown to specifically weaken EAE disease and immune cell infiltration^6,77,78^, the role of CNS parenchymal MHC class I expression in mediating CD8^+^ activation remains unknown. Observations from MS pathological studies document CD8 activation in the CNS. Extensive CD8 clonal expansion to cognate antigen of cells localized to the parenchyma has been observed^15,16,79,80^. Furthermore, CD8s outnumber CD4s in MS brain lesions that are also populated by recruited OPCs^81–83^. Our results, showing that OPCs cross present antigen on MHC class I molecules and activate CD8s, may explain the longstanding pathological observations of CD8 predominance in the MS lesion. We also suggest that OPCs may have this signaling pathway in order to participate in responses to CNS infections. The subsequent OPC death may be an acceptable resolution to acute infection/inflammation given their abundance in the CNS, much in the way dendritic cells die after presenting antigen in the periphery.

Providing context to the contribution of OPC antigen presentation amongst CNS resident cells and known infiltrative APCs is important to better understand MS disease pathogenesis and more generally neuroinflammation. A prior report from one of us (JG) documented that Tip-DCs cross present antigens in EAE^52^. This studied also identified that mature oligodendrocytes, but not microglia were able to mediate determinant spreading and activation of CD8 T-cells. Here we show OPCs express higher levels of peptide loaded class I molecules than do mature oligodendrocytes. Importantly, we were able to demonstrate cross presentation of MBP79-87 MHC class I restricted peptide by OPCs, but similar to the previous study, were unable to detect this peptide loaded MHC class I molecule expression within the microglial population. When comparing OPCs to known APCs such as DCs or macrophages, there are several similarities. As we showed here, OPCs engulf and present peptide with similar kinetics as have been seen in DCs^51^. We observed upregulation of MHC class II and several co-stimulatory molecules in our microarray, in addition to verifying MHC class II surface expression on OPCs (data not shown). The expression of functional MHC class II and co-stimulatory molecule expression is similar to DCs and B-cells; however, further studies need to be performed to further interrogate similarities in their response to cytokine stimulation. Examination of the ability of OPCs to activate CD4^+^ T-cells is clearly of relevance to MS, and would be an important additional future area of study. Furthermore, the relative contribution of OPCs as compared to dendritic cells or other antigen presenting cells would require conditional deletion of part of this pathway, which is a goal of future studies.

Cytotoxicity through Fas/FasL and perforin/granzyme B signaling was another important observed outcome of CD8-MHC class I-OPC interaction^18–23^. This finding raises the question as to why OPCs would contain transcriptional programs that result in their presenting antigen to cytotoxic CD8 T-cells and thereby becoming targets for depletion during periods of IFNγ mediated inflammation. The cross-presentation pathway is thought to exist as a mechanism to facilitate clearance of virally infected cells^49^, and aberrant induction of this pathway could occur in states of chronic inflammation, as occurs in autoimmune diseases in which immune risk genes that regulate transcriptional activation of immune cells may also promote the same immune pathways in glia. Taken together, some OPCs may be aberrantly targeted for cell death in chronic inflammatory diseases of the brain, while others persist and perpetuate bystander activation of immune cells, further propagating the process. In the post-mortem MS plaques, we saw a substantial proportion of OPCs were present and had high PSMB8 expression, the specific immunoproteosome subunit, but only in the areas of failed remyelination and not the normal appearing white matter. This is consistent with the notion that they are either arrested in their maturation and or have taken on an alternative role to present antigen. Since mature oligodendrocytes from MS tissue have been shown to be immunoreactive for Fas surface expression, we propose that OPCs may be targeted for Fas mediated cells death in MS as well^84^. Furthermore, perforin and granzyme signaling pathways are also important for target cytotoxicity by predominantly CD8+ CTLs and natural killer cells. Recent work done by the IMSCG identified a coding variant in PFR1, the gene that encodes perforin, to be associated with the risk of getting MS^85^. This result underlines the potential importance of CD8+ cytotoxicity in MS pathogenesis. Thus, not only do the OPCs fail to differentiate, but they are co-opted by the immune system to propagate the CNS immune response.

If this process were operational in MS, as suggested by our observation of marked expression of PSMB8 only in areas of MS brain demyelination, it could suggest that either depleting or redirecting the transcriptional program of pathogenic OPCs may be necessary for myelin repair. Targeting these newly discovered inflammatory signaling pathways in OPCs may be an important step in blocking inflammatory responses of OPC, and to facilitate therapies designed to promote myelin repair, which to date have targeted developmental pathways. While proteasomal inhibitors are used in malignancies such as myeloma, they are quite toxic. Nevertheless, targeting one of the three critical subunits of the immunoproteasome (PSMB8, 9, and 10) might allow selective suppression of antigen cross presentation without suppressing normal cell function^86^.

In summary, we have found a previously unknown antigen presenting function of OPCs in the inflamed CNS, which is mediated by IFNγ induction of the immunoproteasome. This pathway not only suppresses OPC maturation and therefore remyelination, but also promotes cytotoxic CD8 activation and OPC death. Future studies of inhibitors of this pathway may have dual roles in suppressing OPC inflammatory responses and promoting myelin repair, which are not targeted by present MS therapies and could lead to more effective treatments for progressive forms of MS.

## Methods

### Mice

Mice were housed and maintained in a pathogen-free animal facility at Johns Hopkins University. *TRE/IFNγ* and *GFAP/tTA* mice were a kindly provided by B. Popko (University of Chicago) and were then crossed and maintained in-house. OVA257-264 TCR transgenic, OT-1 mice (C57BL/6-Tg(TcraTcrb)1100Mjb/J), and 2D2 MOG35-55 TCR transgenic mice (C57BL/6-Tg(Tcra2D2,Tcrb2D2)1Kuch/J) and were originally purchased from Jackson Laboratory and maintained in-house. C3HeB/Fej (000658) were purchased from Jackson Laboratory as needed for experimentation. C57BL/6 (556) and SAS SD timed-pregnant rats were purchased from Charles River (SD400) for the purpose of postnatal OPC and glia cultures. *PDGFRα-CRE^ER^* were provided by D. Bergles then backcrossed to the C57BL/6 background for 12 generations before being crossed with *Rosa26-YFP*. All animal protocols were approved and adhered to the guidelines of Johns Hopkins Institutional Animal Care and Use Committee.

### Recombinant Cytokines and Pharmacological Inhibitors

To determine the cytokine effects on OPC differentiation, microarray and qPCR, rat recombinant IFNγ (10 ng/mL; Peprotech) or rat recombinant IL-17 (50 ng/mL; R&D Systems) were supplemented. The antigen presentation studies mouse OPCs were used and cultured for 12 hrs (unless otherwise specified) with mouse recombinant IFNγ (10 ng/mL; Peprotech) in the presence of media supplemented with human recombinant PDGF-AA (20 ng/mL; R&D Systems). Pharamacological blockade of different aspects of the cytosolic MHC class I pathway was done using Chloroquine (100 μM; Sigma Aldrich), ONX-0914 (30 nM; ApexBio), and a Cathepsin S (10 nM; Calbiochem) inhibitor.

### Peptides

*In Vitro Studies*: Peptides or protein used for OPC presentation studies were MOG37-50 (50 μg/mL; Johns Hopkins Synthesis and Sequencing Core), OVA257-264 (50 μg/mL; Johns Hopkins Synthesis and Sequencing Core) or Ovalbumin protein (500 μg/mL; Sigma Aldrich). *In Vivo Studies:* MOG1-125 (100 mg/mouse; MedImmune) was used for C3HEB/Fej immunizations. AT experiments used MOG35-55 or MOG97-114 (20 μg/mL; Johns Hopkins Synthesis and Sequencing Core) to select and reactivate isolated CD4 *ex vivo*.

### Primary Cultures

*OPC:* Cerebral cortices were dissected from P4-P6 rodent pups. To obtain cultures with high purity, OPCs were positively selected using A2B5 magnetic beads (Miltenyi) as previously described^37^. A2B5^+^ cells were plated and expanded for 3-4 days with OPC media supplemented with recombinant human PDGF-AA until optimal density was reached and natural BMP antagonist Noggin to inhibit in vitro OPC to astrocyte differentiation and recombinant human PDGF-AA until optimal density was reached. If OPCs were being used in experimentation cells were maintained in PDGF-AA with the addition of cytokines or peptides. For differentiation assays, 96 hrs of culture combinations of T3 (10 nM; Sigma Aldrich) and recombinant cytokine were supplemented into OPC media.

*Astrocyte and Microglia:* Cerebral cortices were dissected from P4-P6 rat pups were mechanically and enzymatically (2.5% trypsin) digested to obtain a single cell suspension before plating on poly-L lysine (PLL) coated T75 flasks. Mixed glia were cultured in glia medium (DMEM, high glucose with 10% heat-inactivated FBS (Hyclone) and 1% Penicillin/Streptomycin) for 8 to 9 days or until a multi-layer culture of astrocytes, OPCs and microglia was achieved. To remove microglia flasks were shaken at 180 rpm for 1 hour. After shaking supernatant was removed and microglia were plated on PLL coated plates and cultured in microglia medium (2% heat-inactivated FBS). The medium was replenished and OPCs were shaken off for 8 hours at 240 rpm. The supernatant containing OPCs was discarded and astrocytes were removed by trypsin then seeded onto PLL coated plates in glia medium (described above). 24 to 48 hrs after plating microglia and astrocytes IFNγ stimulation followed by OVA257-264 addition was begun (Identical experimental protocol as OPCs).

### CD8 Isolation

OVA peptide-specific CD8s were isolated from 8-12 week old OT-1 transgenic mice. Briefly, a single cell suspension was prepared from spleen and lymph nodes. A CD8 negative selection was performed (Stemcell Technologies) and purified cells were stained with Cell Proliferation Dye eFluor450 (Fisher Scientific) according to the manufacturer’s protocol. After flow analysis using Vβ5, CD8, CD62L, CD44 to test for purity and activation status, CD8s were then cultured with OPCs (see below).

### OPC-CD8 Co-Culture

OPCs were prepared for coculture (see above), then stimulated with IFNγ for 12 hours (unless otherwise specified) to allow for transcription and translation of MHC class I antigen presentation pathway mediators. Next, OVA257-264 or Ovalbumin was spiked into the medium and incubated for 8 to 12 hrs to permit for processing and presenting time. Cultures were washed so that the only source of antigen was already processed and presented by the OPCs. Isolated and stained OT-1 CD8s were plated with OPCs at a 3:1 CD8:OPC ratio in medium that was 50% CRPMI (RPMI 1640 (Invitrogen) + 10% FBS + 10 mM HEPES buffer (Quality Biological) + 1 mM sodium pyruvate (Sigma-Aldrich) and MEM NEAA (Sigma-Aldrich) + βME + penicillin and streptomycin) and 50% OPC medium. Between 24-48 hours of culture, cells were collected and analyzed by flow cytometry (see below).

### Adoptive Transfer – Cuprizone Studies

*MOG35-55 CD4 AT-CPZ into* *PDGFRα-Cre^ER^ x Rosa26-YFP*: PDGFRα-Cre^ER^ bred to the C57BL/6 background were crossed with Rosa26-YFP. At the age of 8-12 weeks crossed mice were transferred to a 0.2% CPZ diet for a total of 4 weeks. After 3 weeks of the CPZ diet, 4-hydroxytamoxifen (1 mg/mouse/day for 3 days) was injected to induce CRE recombination in PDGFRα expressing cells. MOG35-55 specific CD4^+^ T-cells from 2D2 TCR transgenic mice were isolated, purified, polarized to TH17 subtype and expanded *ex vivo* as previously described^35^. The 2D2 CD4 purity was measured and approximately 8-10 million cells were injected, IP, into lineage tracing syngeneic recipient mice at the 4 or 6 week CPZ time point. Simultaneously the recipient mice were placed back on a normal chow diet (**supplementary fig 1/8**). 1-2 weeks after, AT mice were sacrificed and prepared for either IHC or flow cytometry analysis (below). *MOG97-114 AT-CPZ in C3HeB/Fej:* Donor C3HeB/Fej mice were immunized with 100 μg of whole rat MOG1-125 protein in complete Freund’s adjuvant (8 mg/mL heat-killed *Mycobacterium tuberculosis* + incomplete Freund’s adjuvant). 250 ng/mouse of Pertussis toxin was injected IP on the day of immunization and 2 days after immunization. After 9-10 days donor rMOG1-125 immunized mice were sacrificed, and spleen and lymph nodes were collected. A single cell suspension from these tissues was prepared. CD4^+^ T-cell negative selection was completed (StemCell Technologies) and cells were cultured *ex vivo* with irradiated APCs collected from the spleen of non-immunized C3HeB/Fej mice. Briefly, 50 μg/mL MOG97-114 peptide and 10 ng/mL of IL23 to expand MOG-specific T-cell population and polarize these cells to the TH17 subtype was completed. After 3 days cells were collected and ficolled to remove dead cells. Approximately 10 million CD4+ T-cells were adoptively transferred via an IP injection into recipient mice that were previously on a 0.2% CPZ diet for 6 weeks. Mice were monitored for EAE disease and once a score of 3 or greater was reached they were sacrificed and prepared for flow cytometry analysis (below).

### Immunostaining

*ICC*: Performed on cultured OPCs after IFNγ and OVA peptide incubation. Prior to fixation APC-α-H2Kb-OVA (1:100; Biolegend) was added to media after washing the cells. This antibody was incubated with live cells for 2 hours before fixation. After fixation α-rabbit-PDGFRα (1:1000; W. Stallcup; Burnham Institute) was used to verify culture purity and for colocalization purposes. DAPI counterstain was performed for 10 minutes then washed prior to slide mounting.

*IHC*: Mice were IP injected with sodium pentobarbital (250 μL of 5mg/mL) and euthanized via cardiac perfusions with 30 mL PBS followed by 10 mL of 4% PFA. The whole brain was collected, post-fixed in 4% PFA for 12-18 hours, transferred to 30% sucrose for 3 days, cryosectioned (30 μm) and kept in antifreeze, and stored in at −20°C until staining was performed. Black Gold II (Millipore) was used to determine myelin content in the corpus callosum. Sections at approximately −0.5 mm, −1.0 mm and −2.0 mm posterior to bregma were stained with α-mouse-MBP SMI-99 (1:1000; eBioscience) and α-rabbit-CD3 (1:50; GeneTex) to determine myeline staining and lymphocyte infiltration, respectively. Parallel sections were stained with α-chicken-YFP (1:1000; Abcam), α-rabbit-PDGFRα (1:600; W. Stallcup; Burnham Institute) and α-mouse-CC1 (1:50; Chemicon) to determine the fate of recombined OPCs. In addition, α-mouse-Olig2 (1:1000; Millipore) was used in combination with α-chicken-YFP to verify that PDGFRα driven CRE recombination within the corpus callosum was restricted to the oligodendrocyte lineage.

*Human tissue IHC:* All brains were collected as part of the tissue procurement program approved by the Cleveland Clinic Institutional Review Board. Details are presented in Supplementary Tables T1. All control and MS brain tissues were characterized for demyelination by immunostaining using 30 µm fixed tissue sections and proteolipid protein (PLP) as described previously (ref 8 in this paper). Adjacent sections were used for double labeling using PLP/PSMB8 and SOX10/PSMB8. Primary antibodies used were rat anti-proteolipid protein (1:250, gift from Wendy Macklin, University of Colorado, Denver), goat anti-SOX-10 (1:100, R&D Systems, Minneapolis, MN), and rabbit anti-PSMB8 (1:1000, Thermo Fisher Scientific, Rockford, IL). Secondary antibodies were biotinylated donkey anti-rat IgG, donkey anti-goat IgG, and donkey anti-rabbit IgG (1:500, Vector Laboratories, Burlingame, CA), and Alexa 488 donkey anti-rat IgG, Alexa 488 donkey anti-goat IgG and Alexa 594 donkey anti-rabbit IgG (1:500, Invitrogen, Carlsbad, CA). Immunofluorescently-labeled tissues were analyzed using a Leica DM5500 upright microscope (Leica Microsystems, Exton, PA). PLP-PSMB8 and PSMB8-Sox10 images were collected from corresponding sections and manually aligned. Resultant images were analyzed using Fiji version (http://fiji.sc) of the free image processing software ImageJ (NIH, http://rsbweb.nih.gov/ij) to evaluate co-localized PSMB8 and SOX10 positive cells and presented as cells/mm2.

### Flow Cytometry

*In vitro:* OPCs or CD8s cultured in vitro were removed from their culture vessel by repeated pipetting. For staining panels that interrogated cytokine and granular protein production cell stimulation was done for 8 hrs using Cell Stimulation Cocktail with protein transport inhibitors (1:250; eBioscience). Live/Dead staining using LIVE/DEAD Fixable Aqua Dead Cell Stain Kit (Thermofisher) was performed for 20 minutes. Surface staining using the following CD8 T-cell markers; APC-CD8a (1:100; eBioscience), PE-α-Vβ5 (1:100; Biolegend), and APC eFluor 780-α-CD3 (1:100; eBioscience) was completed for 30 minutes. In addition, surface markers for T-cell activation were used; APC eFluor 780-α-CD44 (1:100; eBioscience), APC-α-CD62L (1:100; Biolegend), PerCP Cy5.5-α-CD25 (1:100; Biolegend), and FITC-α-CD69 (1:100; Biolegend). Cell permeabilization and fixation was done using IC Fixation Buffer (ThermoFisher) for 20 minutes. Intracellular staining was performed using FITC, PerCP Cy5.5-α-IFNγ (1:100; Biolegend), PE Cy7-α-TNFα (1:100; Biolegend), APC-α-IL-17 (1:100; Biolegend), FITC-αGM-CSF (1:100; eBioscience), and PE-α-perforin (1:100; Biolegend). Staining panels that did not require intracellular staining were immediately stained with a LIVE/DEAD Fixable Aqua Dead Cell Stain Kit (ThermoFisher) after removal from the culture vessel. Surface staining for OPCs was completed using PE Cy7-α-PDGFRa (Miltenyi; 1:25), APC-α-A2B5 (Miltenyi; 1:25), PerCP Cy5.5-α-CD11b (Biolegend; 1:100), BV-421-α-CD11c (Biolegend; (1:100), PE-α-Fas (eBioscience; 1:100), FITC-α-H2Kb (eBioscience 1:100). Simultaneously staining using Caspase 3/7 Green or Red reagent (Essen Biosciences) was completed for 1 hour. Data represented in diagrams and analysis is negative for CD11b and CD11c and positive for PDGFRα and A2B5. Stained cells were run on an 8 color MACSQuant Analyzer (Miltenyi).

*Ex Vivo*: AT-CPZ mice were IP injected with sodium pentobarbital (250 μL of 5mg/mL), after which a cardiac perfusion with 30 mL of PBS was completed. The whole brain was isolated and finely chopped to prepare for efficient enzymatic digestion of tissue using Neural Tissue Dissociation Kit with papain (Miltenyi) according to the manufacturer’s protocol. The remaining tissue was further processed through a 100 μm pore plastic filter to collect a single cell suspension from each brain sample. The myelin component and debris were removed by a 35% percoll gradient centrifugation for 20 minutes, no brake at 650 x g. The cell pellet was resuspended and used for flow cytometry staining. Live/Dead Aqua staining (Thermofisher) was performed for 20 minutes followed by a mouse CD16/CD32 Fc block (1:50; Biolegend) for 10 minutes. To analyze T-cell infiltration APC-eFluor 780-α-CD3 (1:100; Biolegend), eFluor 450-α-CD4 (1:100; Biolegend), APC-α-CD8 (1:100; Biolegend) were used. Dendritic cell, macrophages and microglia populations were analyzed based on the expression of APC-eFluor 780-α-CD45 (1:100; eBioscience), BV-421,PE-α-CD11c (1:100; Biolegend), and PerCP Cy5.5-α-CD11b (1:100; Biolegend). The OPC population was identified using PE Cy7-α-PDGFRα (1:25; Miltenyi) in combination with APC-α-A2B5 (1:25; Miltenyi) or PE-α-O4 (1:25; Miltenyi) in combination with APC-α-A2B5. General MHC class I presentation staining was done for mice on the C57BL/6 and C3HeB/Fej backgrounds using PE,APC-α-H2Kb (1:100; eBioscience) and FITC,eFluor 450-α-H2Kk (1:100; eBioscience), respectively. MBP79-87 specific H2Kk staining was performed using PE,FITC-αH2Kk-MBP79-87 (J. Goverman/Rockland). Finally, to identify cells undergoing cell death and Fas expression interrogation Caspase 3/7 Green or Red reagent (1:1000; Essen Bioscience) and PE-α-Fas (1:100; Biolegend) were utilized. Stained samples were analyzed on an 8 color MACSQuant Analyzer (Miltenyi).

### Gene Array Analysis

Affymetrix microarrays were completed on control and IFNγ treated OPC cultures at baseline (PDGF D0), 8hrs, 24hrs, 48hrs and 96hrs after IFNγ stimulation was initiated. Gene set enrichment analysis (GSEA) was done to by comparing T3 treated samples to T3 + IFNγ treated samples to determine pathway enrichment for genes associated with IFNγ signaling. Data from GSEA was and used in the targeted heat map (**fig. 2e**).

### Live Cell Imaging

OPCs were stained with an eFluor 633 dye prior to the addition of CD8s to ensure subsequent quantification was restricted to the OPC population. Unstained OT-1 CD8s were isolated and added to the stimulated and stained OPCs. Cell membrane permeable, Caspase 3/7 Green reagent (1:1000; Essen Biosciences) was added to determine caspase activity. The Caspase 3/7 reported was designed to report on caspase 3/7 activity by targeted cleavage of the DEVD peptide sequence and subsequent nuclear translocation of the linked DNA binding dye (Essen Biosciences). Images were captured every 3 hours using IncuCyte S3 Live Cell Analysis System (Sartorius). Cultures were analyzed for Caspase 3/7 stain colocalization with the labeled OPCs and the applied mask was counted as particles per image with 4 images taken. A total of three experiments were completed.

### Quantitative PCR

RNA was extracted from cultured OPCs using RNeasy Plus Mini Kit (Qiagen). First-strand cDNA was synthesized using iScript cDNA Synthesis Kit (Bio-Rad). Quantitative PCR was completed using SensiMix (Bioline) a SYBR based reagent and a CFX384 Touch Real-Time PCR Detection System (Bio-Rad). All target genes were normalized to *hprt1* reference gene and delta-delta CT analysis was performed.

### Statistical Analysis

Unpaired two-tailed Student’s t-test, Pearson Correlation, one-way ANOVA with Tukey’s multiple comparisons *post hoc* test and two-way ANOVA with Tukey’s multiple comparisons *post hoc* test were performed. Specific tests are noted in figure legends with significance level annotation. All error bars represent the standard deviation of the mean.

## Acknowledgements

This research was supported by US National Institute of Health grants NS-R37041435 (to PC), the Miriam and Sheldon Adelson Medical Research Foundation (to DB); a NMSS Collaborative Center Award, The Race to Erase MS, and MedImmune. We would like to thank Dr. Brian Popko (University of Chicago) for kindly providing us with the TRE/IFNγ and GFAP/tTA transgeneic mouse lines and Dr. William Stallcup (Burnham Institute) for kindly providing the α-rabbit-PDGFRα antibody.

## Author Contributions

L.A.K., M.D.S., J.J. and P.A.C. designed experiments. L.A.K. executed and analyzed the majority of experiments described in figures and text (unless otherwise stated). J.J. and M.D.S executed initial experiments with AT-Cup lineage tracing in **fig. 1 and supplemental fig 1-2**. J.G.C. completed quantitative PCR validation of microarray findings in **fig 2 and supplemental fig. 5**. K.A.M. managed mouse husbandry, genotyping and phenotyping. J.W. analyzed microarray and performed GSEA. R.D. provided data and analysis for MS patient tissue staining in **fig. 8**. D.B., J.G., T.D. J.K. provided critical and continual feedback regarding experimentation and data analysis. J.G., T.D., J.K., created and provided important reagents used in experiments. L.A.K and P.A.C. wrote the manuscript.

